# Simultaneous detection of pathogens and antimicrobial resistance genes with the open source, cloud-based, CZ ID pipeline

**DOI:** 10.1101/2024.04.12.589250

**Authors:** Dan Lu, Katrina L. Kalantar, Victoria T. Chu, Abigail L. Glascock, Estella S. Guerrero, Nina Bernick, Xochitl Butcher, Kirsty Ewing, Elizabeth Fahsbender, Olivia Holmes, Erin Hoops, Ann E. Jones, Ryan Lim, Suzette McCanny, Lucia Reynoso, Karyna Rosario, Jennifer Tang, Omar Valenzuela, Peter M. Mourani, Amy J. Pickering, Amogelang R. Raphenya, Brian P. Alcock, Andrew G. McArthur, Charles R. Langelier

## Abstract

Antimicrobial resistant (AMR) pathogens represent urgent threats to human health, and their surveillance is of paramount importance. Metagenomic next generation sequencing (mNGS) has revolutionized such efforts, but remains challenging due to the lack of open-access bioinformatics tools capable of simultaneously analyzing both microbial and AMR gene sequences. To address this need, we developed the Chan Zuckerberg ID (CZ ID) AMR module, an open-access, cloud-based workflow designed to integrate detection of both microbes and AMR genes in mNGS and whole-genome sequencing (WGS) data. It leverages the Comprehensive Antibiotic Resistance Database and associated Resistance Gene Identifier software, and works synergistically with the CZ ID short-read mNGS module to enable broad detection of both microbes and AMR genes. We highlight diverse applications of the AMR module through analysis of both publicly available and newly generated mNGS and WGS data from four clinical cohort studies and an environmental surveillance project. Through genomic investigations of bacterial sepsis and pneumonia cases, hospital outbreaks, and wastewater surveillance data, we gain a deeper understanding of infectious agents and their resistomes, highlighting the value of integrating microbial identification and AMR profiling for both research and public health. We leverage additional functionalities of the CZ ID mNGS platform to couple resistome profiling with the assessment of phylogenetic relationships between nosocomial pathogens, and further demonstrate the potential to capture the longitudinal dynamics of pathogen and AMR genes in hospital acquired bacterial infections. In sum, the new AMR module advances the capabilities of the open-access CZ ID microbial bioinformatics platform by integrating pathogen detection and AMR profiling from mNGS and WGS data. Its development represents a critical step toward democratizing pathogen genomic analysis and supporting collaborative efforts to combat the growing threat of AMR.

## Introduction

Antimicrobial resistance (AMR) is responsible for an estimated 1.27 million global deaths annually^1^, and is on track to cause 10 million deaths a year by 2050, becoming a leading cause of global mortality^2^. Furthermore, the World Health Organization has declared AMR to be one of the top ten global public health threats facing humanity^3^.

A critical step in combating AMR is the development and implementation of new methods and analysis tools for genomic detection and surveillance of AMR microbes with high resolution and throughput^4^. Whole genome sequencing (WGS) of cultured bacterial isolates and direct metagenomic next-generation sequencing (mNGS) of biological and environmental samples have emerged at the forefront of technological advances for AMR surveillance^5^. Several tools and databases have been developed over the past decade to enable the detection of AMR genes from both WGS and mNGS data. These include ResFinder^6^, the Comprehensive Antibiotic Resistance Database (CARD)^7,8^, ARG-ANNOT^9^, SRST2^10^, AMRFinderPlus, the Reference Gene Catalog by NCBI^11^, and others.

Effective surveillance for resistant pathogens requires not only detecting AMR genes, but also detecting their associated microbes. Despite this, each task has traditionally been approached separately in bioinformatics pipelines, with few available tools enabling simultaneous evaluation of both. The Chan Zuckerberg ID (CZ ID) mNGS pipeline, for instance, was developed in 2017 to democratize access to metagenomic data analysis through a free, no-code, cloud-based workflow, but has had limited AMR assessment capabilities^12^.

Realizing the unmet need for, and potential impact of, a single bioinformatics tool integrating the detection of both AMR genes and microbes, we sought to add AMR analysis capabilities to the open-access CZ ID mNGS pipeline. Here we report the development of a new AMR module within the CZ ID web platform, which leverages CARD to support openly-accessible AMR detection and analysis. We demonstrate its utility across both WGS and mNGS data, and in clinical and environmental samples, and demonstrate the value of enriching AMR findings through simultaneous unbiased profiling of microbes.

## Implementation

### AMR gene and variant detection using the CZ ID AMR module

The AMR module is incorporated into the CZ ID web application (https://czid.org)^12^ and allows researchers to upload FASTQ files from both mNGS and WGS short-read data. Once uploaded, the module automatically processes samples in the cloud using Amazon Web Services infrastructure, eliminating the need for users to download and install software or maintain high-performance computing resources. The web-based application makes analysis of AMR datasets accessible even to researchers with limited bioinformatics or computational expertise.

Underlying the AMR module is CARD (https://card.mcmaster.ca), a comprehensive, continually curated, database of AMR genes and their variants, linked to gene family, resistance mechanism, and drug class information^7,8^. The AMR module specifically leverages the CARD Resistance Gene Identifier (RGI) tool (https://github.com/arpcard/rgi)^7,13^ to match short reads or contigs to AMR gene reference sequences in the CARD database, returning metrics such as gene coverage and percent identity. CARD also contains a Resistomes & Variants database of *in silico* predictions of allelic variants and AMR gene homologs in pathogens of public health significance. This database provides information linking AMR genes to specific species, and can be used for k-mer-based pathogen-of-origin prediction, a beta feature implemented in RGI^13^.

The CZ ID AMR module automates the running of a containerized WDL workflow that strings together multiple steps and informatics tools to enable efficient data processing and accurate resistome profiling. The workflow shares the same preprocessing steps as the existing CZ ID mNGS module. Briefly, it accepts raw FASTQ files from short-read mNGS or WGS samples as input (DNA or RNA) (**Fig. 1, Fig. S1**). Low quality and low complexity reads are first removed with fastp^14^, host reads are removed with Bowtie2^15^ and HISAT2^16^, and then duplicate reads are filtered out using CZID-dedup (https://github.com/chanzuckerberg/czid-dedup). The resulting quality- and host-filtered reads are subsampled to 1 million single-end reads or 2 million paired-end reads to limit the resources required for compute-intensive downstream alignment steps. In the AMR workflow, to accommodate targeted mNGS protocols designed to amplify many copies of low abundance AMR genes, duplicate reads are then added back prior to further processing.

**Figure 1:**
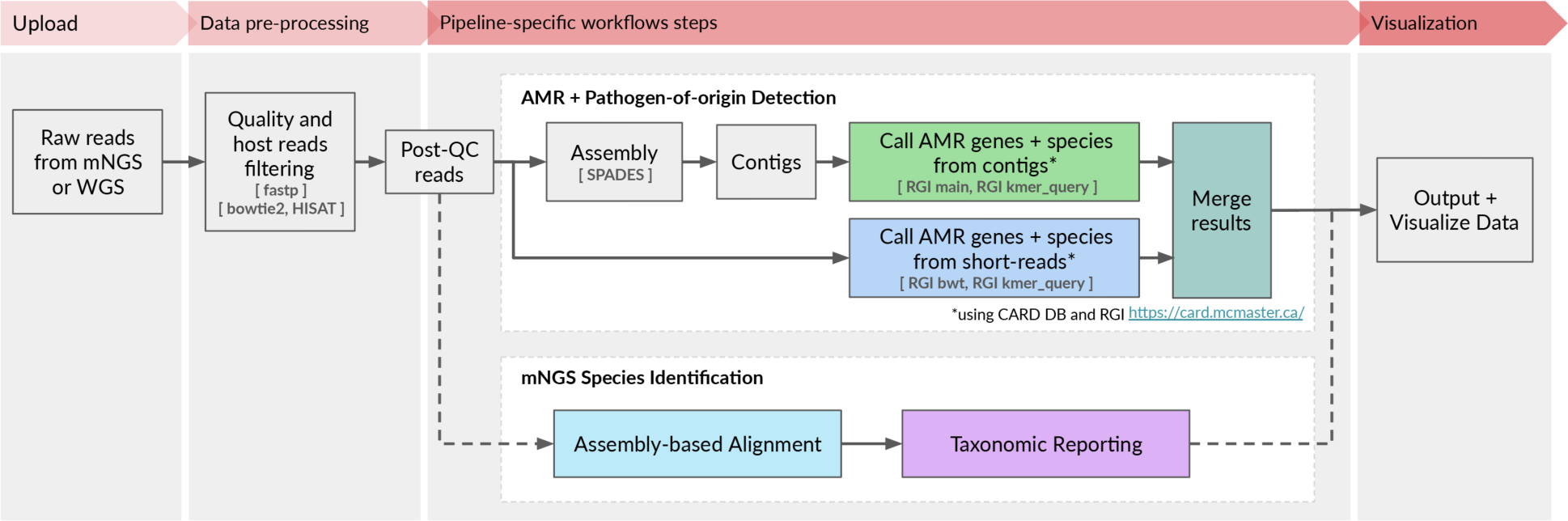
High-level flow diagram highlighting the integrated AMR and mNGS modules within the CZ ID pipeline. A more detailed diagram is provided in Figure S1.

There are two parallel approaches for AMR gene detection (**Fig. 1, Fig. S1**). In the ‘contig’ approach, the short reads are assembled into contiguous sequences (contigs) using SPADES^17^, and the contigs are subsequently sent to RGI (main) for AMR gene detection based on sequence similarity and mutation mapping. In the ‘read’ approach, the short reads are directly sent to RGI (bwt) for read mapping by KMA^18^ to CARD reference sequences (**Fig. 1**). In both approaches, the assembled contigs or reads containing AMR genes are also sent to RGI (kmer_query) for pathogen-of-origin detection.

### AMR module result output

The AMR module displays results in an interactive table, facilitating viewing, sorting, and filtering. The table is organized in three collapsible vertical sections: 1) general Information, 2) contigs, and 3) reads (**Fig. 2A**). The general information section includes “Gene” and “Gene Family” as well as information on the antibiotic(s) against which the gene confers resistance (“Drug Class” and “High-level Drug Class”), resistance mechanism (“Mechanism”), and model used to identify resistance (“Model”). With respect to the latter, several models are used to identify resistance such as the CARD *protein homolog model* which identifies the presence of AMR genes, and the *protein variant model* which identifies specific mutations that confer resistance. Clicking on the AMR gene name will reveal a description and web hyperlinks to CARD, NCBI and PubMed entries.

**Figure 2:**
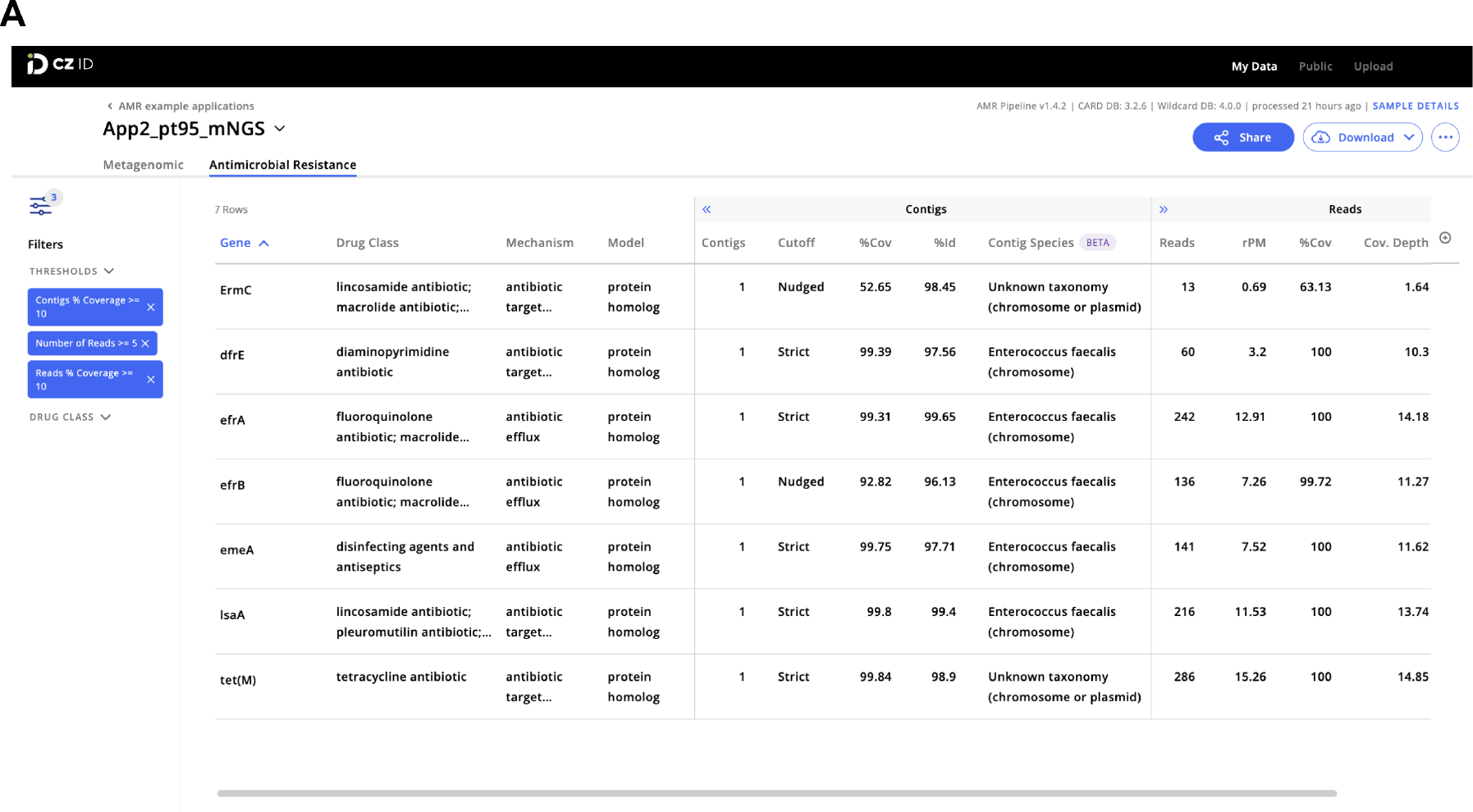
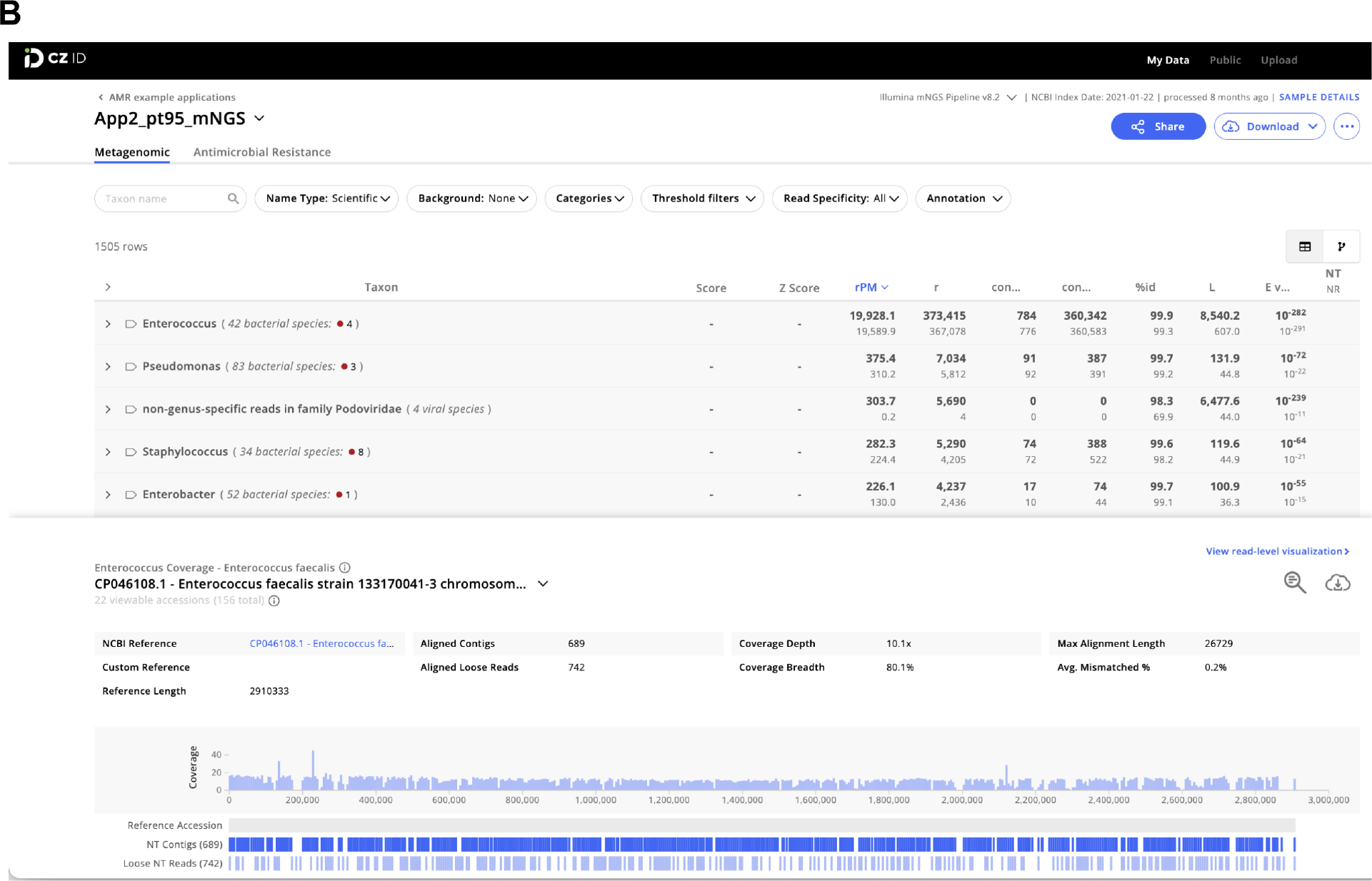
Examples of CZ ID web tool sample reports. **(A)** The report in the AMR module with a filter of Number of Reads >= 5 and Reads/Contig % coverage >= 10% applied to the AMR genes. **(B)** The report in the mNGS module showing the list of detected species and the coverage visualization for one species. Details about report metrics are discussed in the main text and CZ ID help center https://help.czid.org/.

The “Contigs” section includes the number of contigs that map to each AMR gene (“Contigs”), cutoff based on BLAST bit-score (“Cutoff”), percentage of the AMR gene covered by all contigs (“%Cov”), percent identity of the covered region (“%Id”), and pathogen-of-origin prediction based on contigs (“Contig Species”). The “Reads” section includes metrics corresponding to the number of reads mapping to the AMR gene (“Reads”), relative abundance of the AMR gene in reads per million reads sequenced (“rpM”), percentage of AMR gene covered by sequencing reads (“%Cov”), average depth of reads aligned across the gene (“Cov. Depth”), average depth of reads aligned across the gene per million reads sequenced (“dpM”), and a pathogen-of-origin prediction based on reads (“Read Species”). All columns can be sorted and numerical metrics can be further filtered using user specified thresholds.

Results files at each stage of the pipeline can be downloaded for inspection or additional downstream analysis. These files include quality- and host-filtered reads, assembled contigs, AMR annotations and corresponding metrics in tabular format, and all output files from CARD RGI. The contigs as well as short reads mapped to each AMR gene can also be downloaded. The AMR module does not provide heatmap plotting functionality at the moment but users can download the results and use CZ ID’s public scripts to generate heatmaps: https://github.com/chanzuckerberg/czid-amr-heatmap

### Quality filtering for AMR gene predictions

One challenge with mNGS-based AMR surveillance is interpretation of results. The CZ ID AMR module provides key quantitative metrics including rpM, percent coverage of the AMR gene, and dpM to enable assessments of relative abundance and the confidence of AMR gene assignments. Additionally, for AMR detection using contigs, the “Cutoff” column which reports RGI’s stringency thresholds based on CARD’s curated bit-score cut-offs can provide valuable insight into AMR gene alignment confidence. Here, “Perfect” indicates perfect or identical matches to the curated reference sequences and mutations in CARD while “Strict” indicates matches to variants of known AMR genes, including a secondary screen for key mutations.

Finally, the terminology “Nudged’’ is adopted by the CZ ID module to indicate more distant homologs (matched via RGI’s “Loose” paradigm) with at least 95% identity to known AMR genes, which is ideal for discovery but is more likely to return false-positive hits. Given that a consensus approach has yet to be developed for quantifying and interpreting AMR genes from mNGS and WGS data, the CZ ID AMR module provides comprehensive information that can be subsequently filtered or otherwise optimized based on the goals of a given analysis.

### Microbial profiling using the CZ ID mNGS module

The CZ ID mNGS module, which has undergone several updates since first described^12^, preprocesses the uploaded reads and then proceeds to assembly-based alignment to produce taxonomic relative abundance profiles for each sample. Briefly, the non-host reads output by the quality- and host-filtering steps (as described above) are aligned to the NCBI nucleotide (NT) and protein (NR) databases using minimap2^19^ and DIAMOND^20^, respectively, to identify putative short-read alignments (**Fig. 1, Fig. S1**). Then, reads are assembled into contigs using SPADES^17^ and contigs are re-aligned to the set of putative accessions using BLAST^21^ to improve specificity. Finally, alignments are used to identify taxons of origin, which are tallied into relative abundance estimates^12^. The web interface provides various reports with metrics including reads per million (“rpM”), number of reads (“r”), number of contigs (“contig”), number of reads in the contigs (“contig r”), percent identity (“%id”), and average length of alignment (“L”), alongside visualizations and download options to support the analysis and exploration of results (**Fig. 2B**).

### Connecting the pathogens and AMR genes

The CZ ID platform enables simultaneous data analysis of microbe and AMR genes from a single data upload via the mNGS and AMR modules. This provides complementary, but distinct, microbial and AMR gene profiles from a given sample or dataset. The mNGS module does not provide any direct link between species calls and AMR genes from the AMR module, although in cases where a single bacterial pathogen comprises the majority of reads in a metagenomic sample, this may be inferred.

Conversely, the AMR module provides two ways to help connect AMR genes to their source microbes. First, each AMR gene returned in the report table is hyperlinked to its corresponding CARD webpage, where the Resistomes section reports all species in which the gene and its variants have been identified as predicted by RGI. Secondly, the AMR module returns results from a pathogen-of-origin analysis conducted by RGI^13^, which maps k-mers derived from reads or contigs containing the AMR gene of interest against AMR alleles in CARD Resistomes & Variants database. This second approach is particularly useful for identifying the source species in cases when the first CARD Resistomes section lists multiple species or genera. However, because only AMR gene sequences present in CARD are considered in the pathogen-of-origin analysis, as opposed to species identification using complete reference genome sequences in the mNGS module, species predictions from AMR module are best interpreted in the context of all outputs from the CZ ID AMR and mNGS modules.

### Sharing results for collaboration

Projects on CZ ID can be shared with specific users or made public to all users. Everyone with access to the project can view or download the results, and perform data filtering or other analyses. All data and results for this paper can be accessed by searching for a project named “AMR example applications” among public projects at https:///czid.org.

## Results

### Application 1: Identification of AMR genes from WGS and mNGS data

To demonstrate the CZ ID AMR module’s utility for detecting bacterial pathogens and their AMR genes in both WGS and mNGS data, we leveraged data from a recent investigation of transfusion-related sepsis^22^. In this study, two immunocompromised patients received platelet units originating from a single donor. Both developed septic shock within hours after the transfusion, with blood cultures from Patient 1, who did not survive, returning positive for *Klebsiella pneumoniae.* Patient 2, who was receiving prophylactic antibiotic therapy at the time of the transfusion, survived, but had negative blood cultures. Direct mNGS of post-transfusion blood samples from both patients revealed a large increase in reads mapping to *Klebsiella pneumoniae*, a pathogen which was later also identified from culture of residual material from the transfused platelet bag (**Fig. 3A**)^22^. While blood mNGS data yielded less coverage of the *K. pneumoniae* genome compared to WGS of the cultured isolates, mNGS of patient 1’s post-transfusion plasma sample recovered all the AMR genes found by WGS of cultured isolates (**Fig. 3B**). Even in patient 2, whose blood sample had fewer reads mapping to *K. pneumoniae,* most AMR genes found in the cultured isolates were still able to be identified using the RGI “Nudged” threshold.

**Figure 3:**
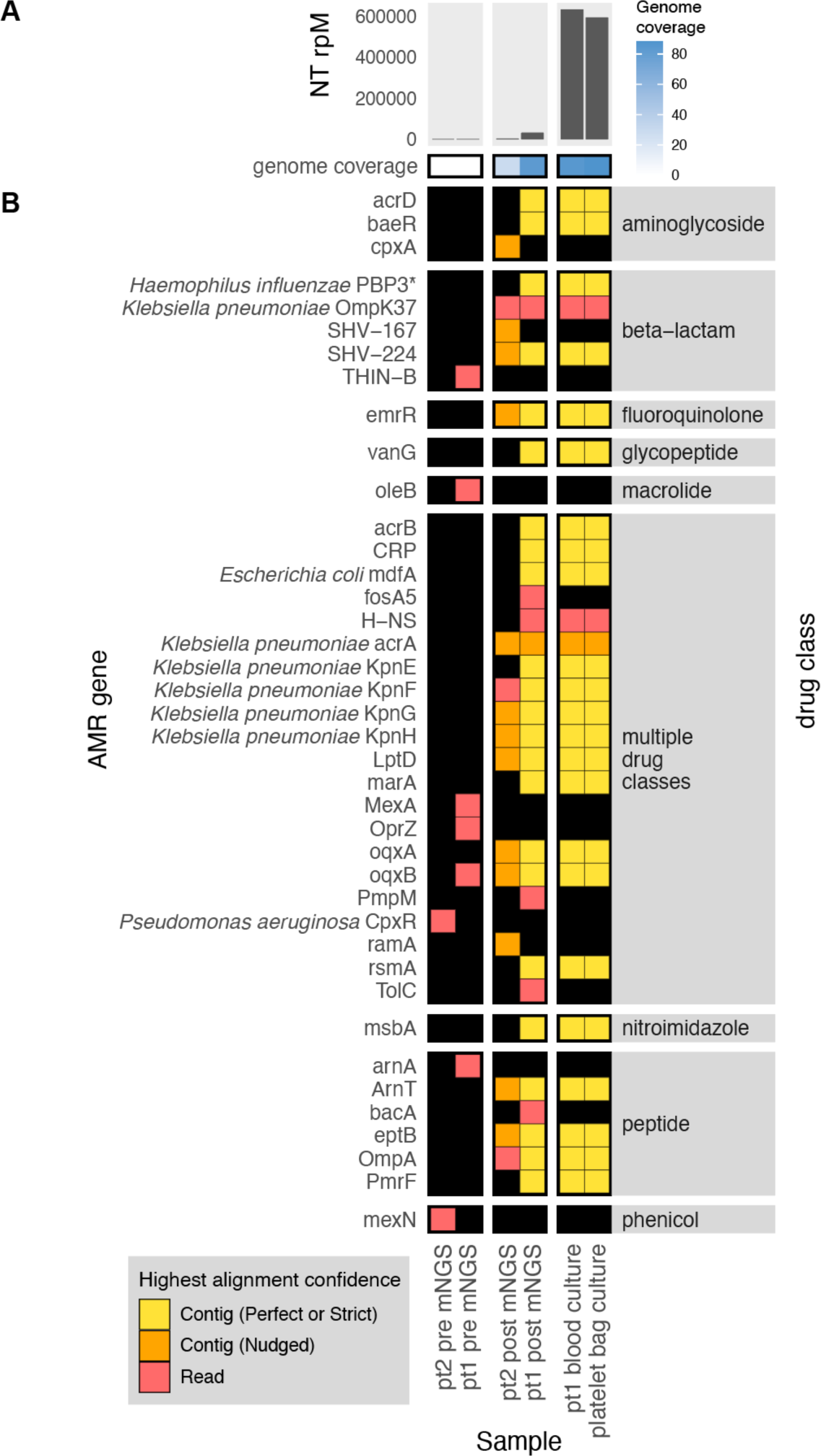
Combining pathogen detection and AMR gene profiling of mNGS and WGS data to investigate *Klebsiella pneumoniae* transfusion-related sepsis. **(A)** Abundance and genome coverage of *Klebsiella pneumoniae* from direct mNGS of plasma or serum samples versus WGS of cultured bacterial isolates. **(B)** AMR genes detected in each sample. *denotes AMR gene(s) for which resistance originates due to point mutations (as opposed to presence/absence of the gene); these were detected by the “protein variant model” in CARD and the gene name shown is a representative reference gene containing the mutations known to lead to resistance. Legend: NT rPM = reads mapping to pathogen in the NCBI NT database per million reads sequenced. Contig = contiguous sequence. Strict/Perfect/Nudged refers to RGI’s alignment stringency threshold. “pt1” = patient 1, “pt2” = patient 2. “pre” = pre-transfusion, “post” = post-transfusion.

### Application 2: Comprehensive metagenomic and WGS profiling of pathogens and AMR genes in the setting of a hospital outbreak

To demonstrate how the CZ ID AMR module can facilitate deeper insights into pathogen and AMR transmission in hospitals, we evaluated WGS and mNGS data from surveillance skin swabs collected from 40 babies in a neonatal intensive care unit (NICU). The swabs were collected to evaluate for suspected transmission of methicillin-susceptible *Staphylococcus aureus* (MSSA) between patients. WGS of the MSSA isolates followed by implementation of the AMR module demonstrated many shared AMR genes, and revealed a cluster of nine samples with identical AMR profiles (**Fig. 4A**). Subsequent phylogenetic assessment using split k-mer analysis with SKA2^23^, revealed that samples within this cluster differed by less than 11 single nucleotide polymorphisms (SNP) across their genomes, consistent with an outbreak involving *S. aureus* transmission between patients (**Fig. 4B**).

**Figure 4:**
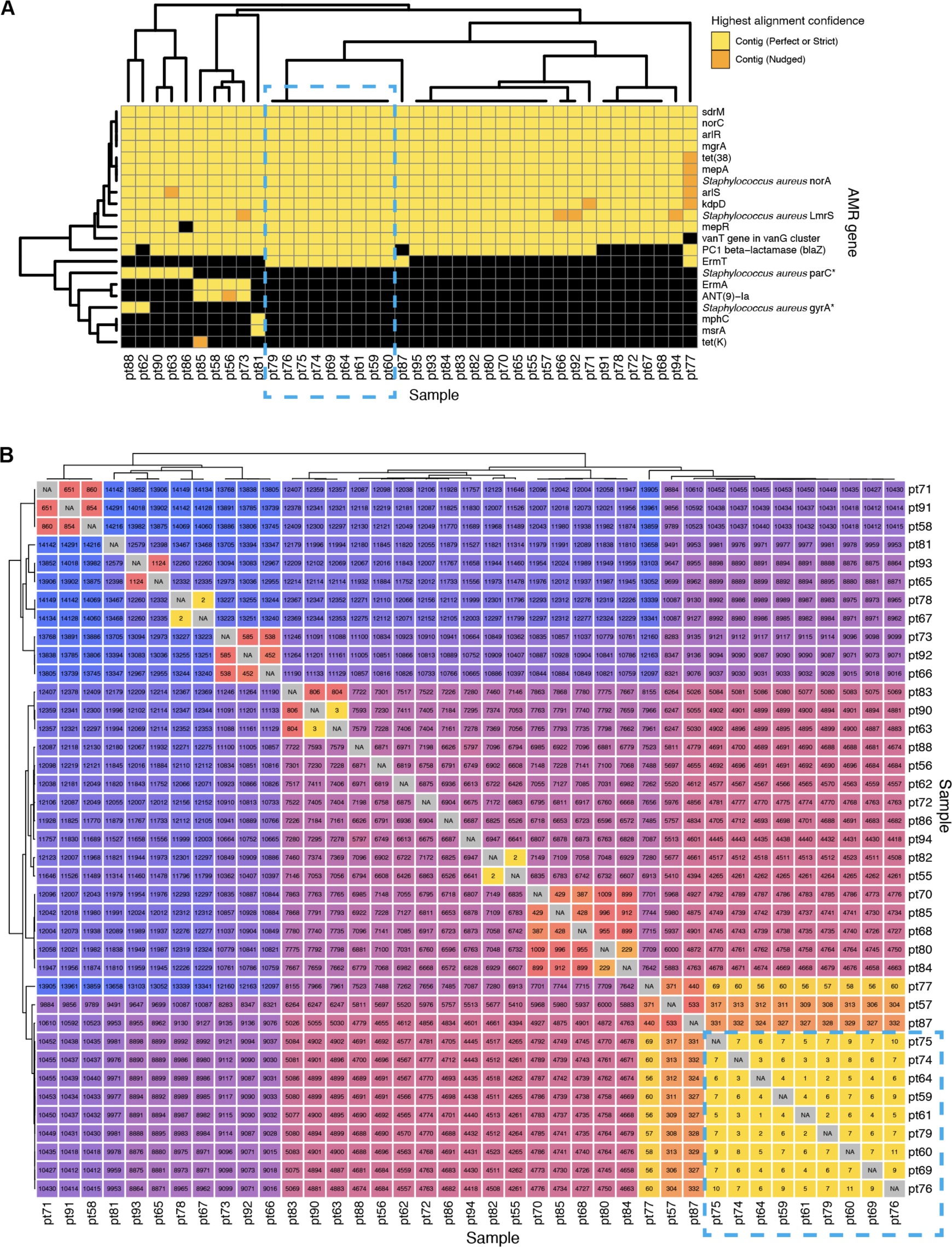
Outbreak investigation pairing WGS of methicillin susceptible *Staphylococcus aureus* isolates and mNGS of surveillance skin swabs from babies in a neonatal intensive care unit. **(A)** Unsupervised clustering of AMR gene profiles from WGS data reveals a cluster of related isolates indicated by the dashed-line box. **(B)** Matrix of single nucleotide polymorphism (SNP) distances between each sequenced isolate confirms the genetic relatedness of this cluster, which is highlighted by a dashed-line box.

Within this cluster of patients, we considered whether other bacterial species in the microbiome were also being exchanged in addition to the *S. aureus*. Intriguingly, mNGS analysis of the direct swab samples from which the *S. aureus* isolates were selectively cultured revealed a diversity of bacterial taxa, many of which were more abundant than *S. aureus*. These included several established healthcare-associated pathogens that were never identified using the selective culture-based approach, such as *Enterobacter*, *Citrobacter*, *Klebsiella* and *Enterococcus* species. mNGS also demonstrated that each sample had a distinct microbial community composition even among samples from the cluster, indicating that only *S. aureus* and potentially a subset of other species were actually exchanged between babies, rather than the entire skin microbiome (**Fig. 5A**).

**Figure 5:**
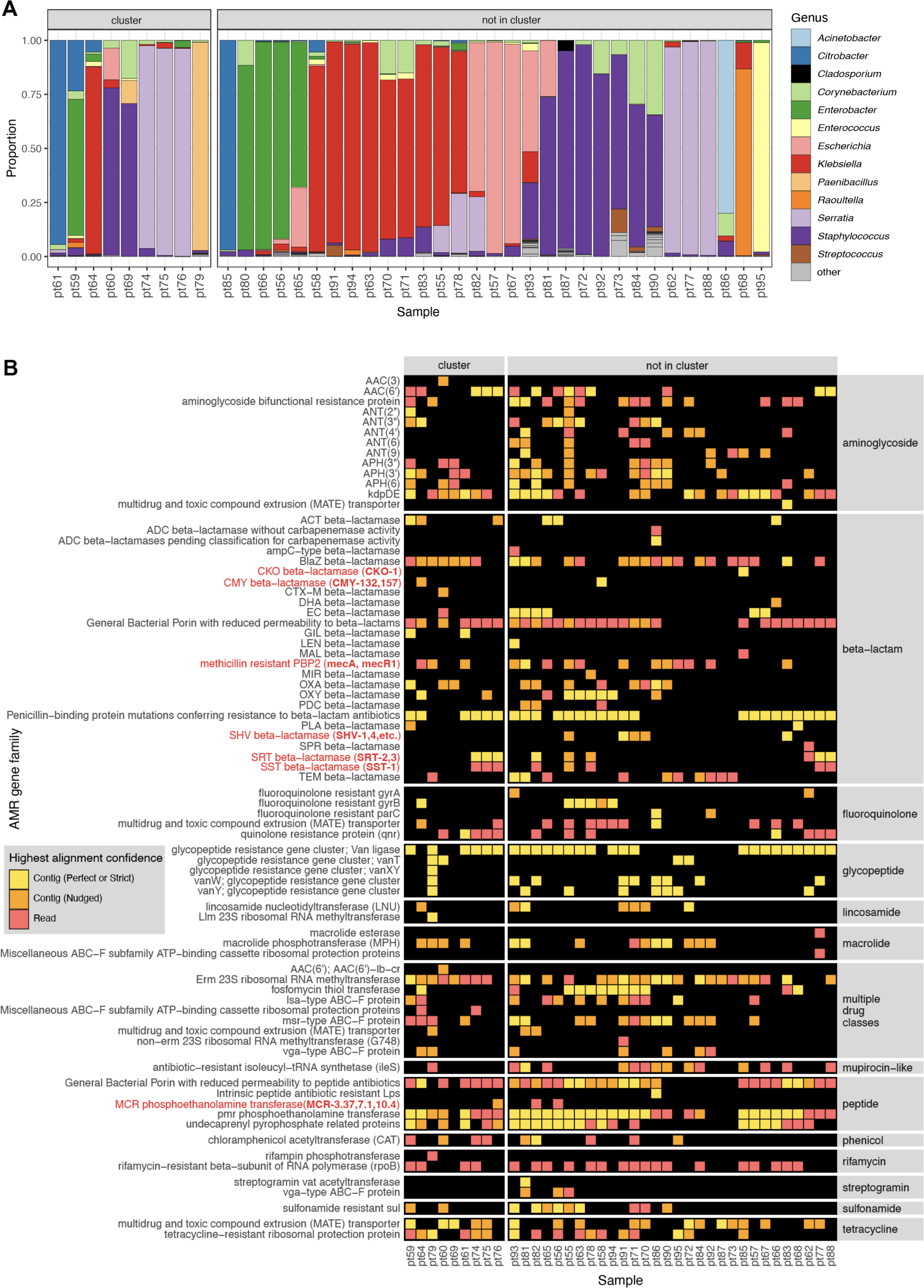
Bacterial genera and AMR genes detected by mNGS of skin swabs from babies in a neonatal intensive care unit. **(A)** mNGS of swab samples demonstrated a diversity of genera in both samples from patients within an outbreak cluster of genetically related *S. aureus*, as well as in those from patients outside of the cluster. **(B)** mNGS analysis revealed a greater number and type of AMR gene families versus those identified by WGS of *S. aureus* isolated in culture from the swabs. Selected AMR gene families of high public health concern are highlighted in red with the specific genes detected in parenthesis.

Further analysis of mNGS data using the AMR module also revealed a diversity of AMR genes conferring resistance to several drug classes, and commonly associated with nosocomial pathogens. These included genes encoding ampC-type inducible beta-lactamases (e.g., *CKO, CMY, SS*T), extended spectrum beta-lactamases (e.g., *SHV*), and the recently emerged *MCR* class of AMR genes, which confer plasmid-transmissible colistin resistance^24^.

The AMR gene profiles varied greatly across the samples, both within the cluster and outside of the cluster, consistent with the observed taxonomic diversity (**Fig. 5B**). Together, these results revealed both inter-patient MSSA transmission in the NICU, and the acquisition of AMR genes associated with nosocomial pathogens within the first months of life.

### Application 3: Correlating pathogen identification with AMR gene detection

Next, we aimed to integrate results from the CZ ID mNGS and AMR modules by analyzing mNGS data from critically ill patients with bacterial infections. In Patient 350^25^, who was hospitalized for *Serratia marcescens* pneumonia, metagenomic RNA sequencing (RNA-seq) of a lower respiratory tract sample identified *Serratia marcescens* as the single most dominant species within the lung microbiome (**Fig. 6A**)^25^. Among the detected AMR genes, based on the Resistomes & Variants information from CARD, *SRT-2* and *SST-1* are found exclusively in *Serratia marcescens* (**Fig. 6B** in blue). Further analysis by the pathogen-of-origin feature in the AMR module matched the k-mers from reads and contigs containing *rsmA, AAC(6’)-Ic,* and *CRP* to *Serratia marcescens* (**Fig. 6B** in purple).

**Figure 6:**
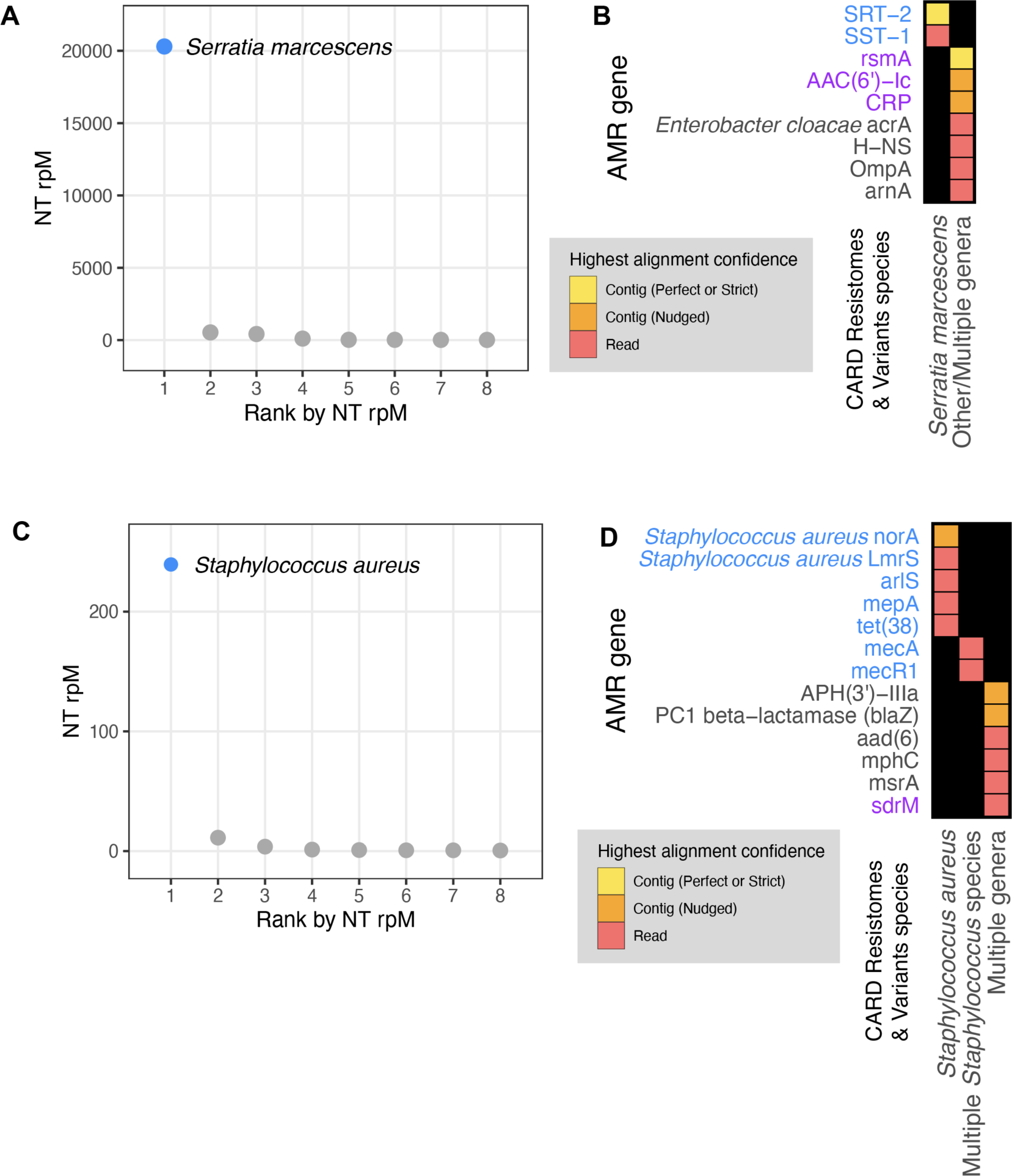
Co-detection of microbes and AMR genes in patients with critical bacterial infections using the CZ ID mNGS and AMR modules. **(A)** Relative abundance (reads per million, rpM) of the eight most abundant taxa in the lower respiratory tract detected by RNA mNGS of tracheal aspirate from a patient with *Serratia marcescens* pneumonia. The dominant microbe is highlighted in blue. **(B)** AMR genes and their species prediction by the AMR module. Columns indicate the species these AMR genes and their variants are found in according to CARD Resistomes & Variants database, and those found in the dominant species as in (A) are colored in blue. AMR genes that are further associated with the dominant species by the pathogen-of-origin analysis are colored in purple. **(C)** Relative abundance (rpM) of the eight most abundant taxa detected by plasma DNA mNGS in a patient with sepsis due to *MRSA* bloodstream infection. The dominant microbe is highlighted in blue. **(D)** AMR genes and their species prediction by the AMR module. Columns indicate the species these AMR genes and their variants are found in according to CARD Resistomes & Variants database, and those found in the dominant species as in (C) are colored in blue. AMR genes that are further associated with the dominant species by the pathogen-of-origin analysis are colored in purple.

In Patient 11827^26^, who was hospitalized for sepsis due to a methicillin-resistant *Staphylococcus aureus* (MRSA) blood stream infection, analysis of plasma mNGS data demonstrated that *Staphylococcus aureus* was the dominant species present in the blood sample (**Fig. 6C**)^26^. Among the detected AMR genes, based on Resistome & Variants information from CARD, *Staphylococcus aureus norA, Staphylococcus aureus LmrS, arlS, mepA, tet(38), mecR1, mecA* are found exclusively in staph species (**Fig. 6D** in blue). Pathogen-of-origin analysis further matched k-mers from the reads containing *sdrM* to *S. aureus* (**Fig. 6D** in purple).

### Application 4: Profiling the longitudinal dynamics of pathogens and AMR genes

To demonstrate the utility of the CZID mNGS and AMR modules for studying the longitudinal dynamics of infection, we analyzed serially-collected lower respiratory RNA-seq data from a critically ill patient with respiratory syncytial virus (RSV) infection who subsequently developed ventilator-associated pneumonia (VAP) due to *Pseudomonas aeruginosa*^27,28^. Analysis of microbial mNGS data using the CZ ID pipeline highlighted the temporal dynamics of RSV abundance, which decreased over time. Following viral clearance, we noted an increase in reads mapping to *P. aeruginosa* on day 9, correlating with a subsequent clinical diagnosis of VAP and bacterial culture positivity (**Fig. 7A**)^27,28^. Analysis using the CZ ID AMR module demonstrated that *P. aeruginosa*-associated AMR genes were also detected, and their prevalence tracked with the relative abundance of the nosocomial bacterial pathogen (**Fig. 7B**).

**Figure 7:**
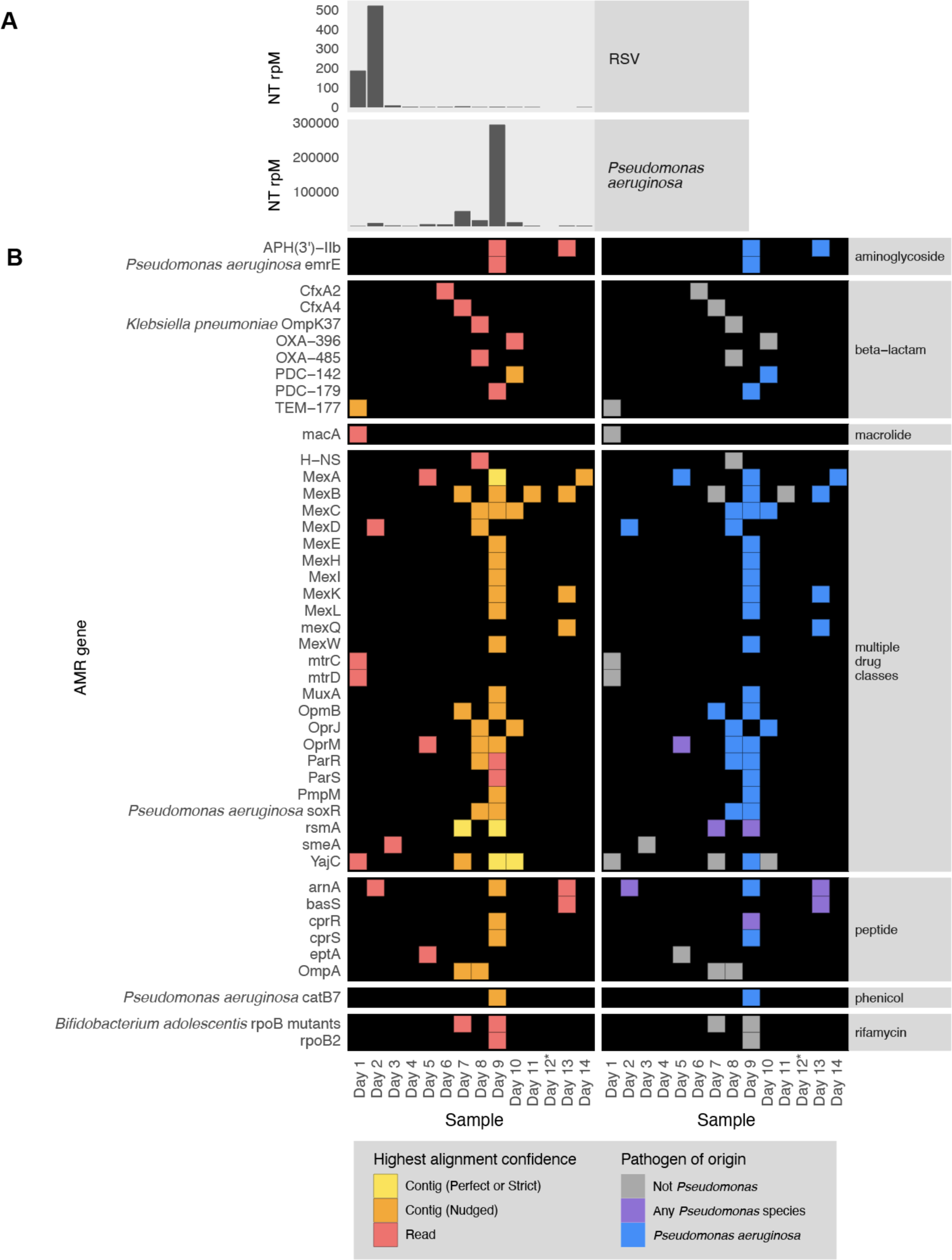
Longitudinal profiling of pathogen and AMR gene abundance in a patient hospitalized for severe Respiratory Syncytial Virus (RSV) infection who developed *Pseudomonas aeruginosa* Ventilator Associated Pneumonia (VAP). **(A)** Relative abundance in reads per million (rpM) of RSV and *P. aeruginosa* detected by the CZ ID mNGS pipeline. **(B)** AMR genes detected in the lower respiratory tract microbiome at each time point. Perfect or strict AMR alignments from contigs are highlighted in yellow, while those nudged are orange. Short read alignments are in red. AMR genes mapping to *Pseudomonas aeruginosa* or any *Pseudomonas* species are highlighted in blue and purple, respectively. *Sample from Day 12 did not have enough sequencing reads but was plotted to maintain even scaling on the x-axis.

### Application 5: AMR gene detection from environmental surveillance samples

Lastly, to highlight the application of the CZ ID AMR module for environmental surveillance of AMR pathogens, we analyzed publicly-available short-read mNGS data from a wastewater surveillance study comparing Boston, USA to Vellore, India^29^. In this study, municipal wastewater, hospital wastewater, and surface water samples were collected from each city and underwent DNA mNGS. From AMR gene alignments at the contig level, we observed a total 22 AMR gene families in Boston samples versus 30 from Vellore (**Fig. 8**). Several AMR genes of high public health concern such as the *KPC* and *NDM* plasmid-transmissible carbapenemase genes were only present in hospital effluent, reflecting the fact that hospitals frequently serve as reservoirs of AMR pathogens^30^.

**Figure 8.**
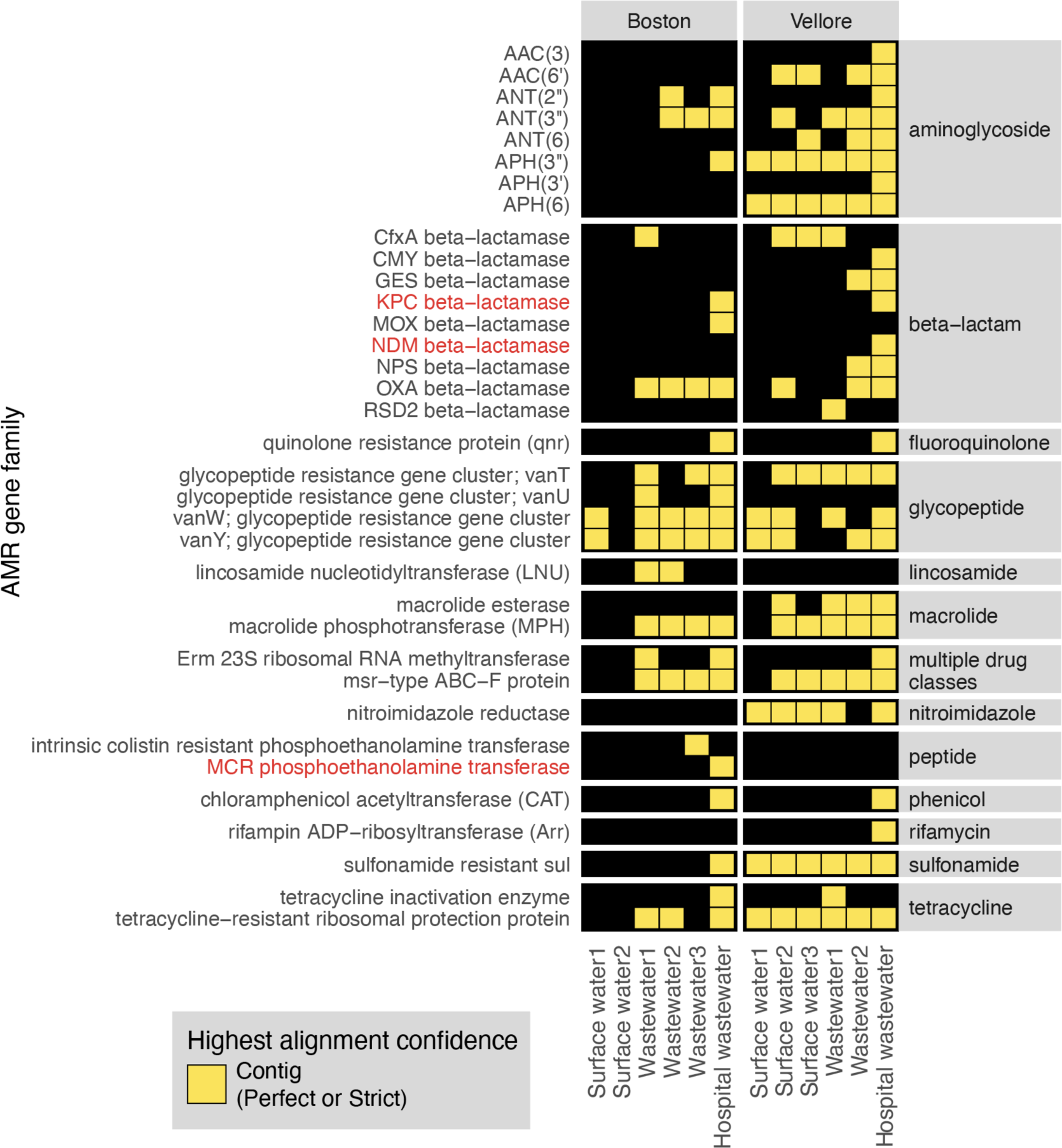
AMR surveillance from environmental water samples. AMR gene families identified from global surveillance of surface or wastewater samples from Boston, USA and Vellore, India. AMR genes found by contigs that passed Perfect or Strict cutoff are included in heatmap. Gene families of high public health concern are highlighted in red.

## Discussion

Metagenomics has emerged as a powerful tool for studying and tracking AMR pathogens in a range of research and public health contexts. Both surveillance and research applications of mNGS benefit from simultaneous assessment of AMR genes and their associated microbes, yet traditionally separate bioinformatics workflows and resource-intense computational infrastructure have been required for each. Here, we address these challenges with the CZ ID AMR module, a fast and openly accessible platform for combined analysis of AMR genes and microbial genomes that couples the expansive database and advanced RGI software of CARD with the unbiased microbial detection capacity of CZ ID. We demonstrate the AMR module’s diverse applications from infectious disease research to environmental monitoring through a series of case studies leveraging four observational patient cohorts and a wastewater surveillance study.

The CZ ID AMR module is designed to enable rapid and accessible data processing without a need for coding expertise, and return a comprehensive set of AMR gene alignment metrics to aid in data interpretation. Researchers can then apply stringency threshold filters to maximize sensitivity or specificity depending on the use case. For instance, when seeking to detect established AMR genes from data types with high coverage of microbial genomes (e.g., WGS data of cultured isolates), “Perfect” or “Strict” stringency thresholds maximize the accuracy of assignments. In contrast, from mNGS data with sparse microbial genome coverage (e.g., from blood or wastewater), using “Nudged” to increase sensitivity of mapping reads at the expense of specificity may be the only way to detect biologically important AMR genes. The “Nudged” threshold also enables more alignment permissiveness to sequence variations, which can be helpful for detecting novel alleles. The CZ ID AMR module provides various metrics to support optimization of cutoffs based on specific sample types and applications by the users.

Depending on the number of reads, breadth of coverage, and whether reads originate from conserved versus variable gene regions, the confidence of AMR gene assignment can vary. Generally, the confidence of contig-based AMR gene assignments is greater than read-based AMR gene matches due to the increased length of assembled fragments. When it comes to AMR gene alleles with high sequence similarity, such as those from within the same gene family, the AMR module can only distinguish between them if sufficient gene coverage is achieved. In most of our analyses, if genes within the same family were identified at both the individual read and contig level, we preferentially evaluated the contig annotation to maximize allele specificity.

As our understanding of AMR gene biology increases over time, annotations may change in the CARD reference database that underpins the CZ ID AMR gene module. This was evident, for instance, in the *Klebsiella* transfusion-related sepsis case (Application 1, **Fig. 2B**), where *mdfA* was annotated as conferring resistance to tetracycline antibiotics based on CARD version 3.2.6, used for our analysis. This will be updated as a multiple drug resistance gene^31^ in the next CARD release. To mitigate database limitations and ensure traceability of results over time, CZ ID periodically updates the database versions and highlights the specific versions of the underlying databases used for each analysis.

CZ ID enables simultaneous detection of pathogens and AMR genes, and our results emphasize the importance of integrating taxonomic abundance from the CZ ID mNGS module with several data outputs within the AMR module. Each AMR gene is directly linked to its CARD webpage where the Resistomes section provides information on the species predicted to harbor the gene of interest and its variants. The pathogen-of-origin predictions, while still a beta feature, can further help identify the source species of detected AMR genes. These assignments are predictions based on matching AMR sequences in each sample to CARD Resistomes & Variants database, and should be interpreted in the context of the microbes found to exist in the sample from the CZ ID mNGS module output. Connecting AMR genes to their originating microbes thus necessitates integrating all available results from both the CZ ID AMR and mNGS modules.

In sum, we describe the novel AMR analysis module within the CZ ID bioinformatics web platform designed to facilitate integrated analyses of AMR genes and microbes. This open-access, cloud-based pipeline permits studying AMR genes and microbes together across a broad range of applications, ranging from infectious diseases to environmental surveillance. By overcoming the significant computing infrastructure and technical expertise typically required for mNGS data processing, this tool aims to democratize the analysis of microbial genomes and metagenomes across humans, animals, and the environment.

## Methods

### Patient enrollment, sample collection and ethics

Skin swabs and cultured isolates analyzed for Application 2 (hospital outbreak) were collected under the University of California San Francisco Institutional Review Board (IRB) protocol no. 17-24056, which granted a waiver of consent for their collection, as part of a larger ongoing surveillance study of patients with healthcare-associated infections.

Samples analyzed for Application 4 (longitudinal profiling) were collected from patients enrolled in a prospective cohort study of mechanically ventilated children admitted to eight intensive care units in the National Institute of Child Health and Human Development’s Collaborative Pediatric Critical Care Research Network (CPCCRN) from February 2015 to December 2017. The original cohort study was approved by the Collaborative Pediatric Critical Care Research IRB at the University of Utah (protocol no. 00088656). Details regarding enrollment and consent have previously been described ^27,28^. Briefly, children aged 31 days to 18 years who were expected to require mechanical ventilation via endotracheal tube for at least 72 hours were enrolled. Parents or other legal guardians of eligible patients were approached for consent by study-trained staff as soon as possible after intubation. Waiver of consent was granted for TA samples to be obtained from standard-of-care suctioning of the endotracheal tube until the parents or guardians could be approached for informed consent.

For all other applications and analyses, previously published datasets were used as described in the data and code availability section.

### Nucleic acid extraction and Illumina sequencing

For the skin swab samples and cultured isolates described in Application 2, DNA was extracted using the Zymo pathogen magbead kit (Zymo Research) according to manufacturer’s instructions. Sequencing libraries were then prepared from 20ng of input DNA using the NEBNext Ultra-II DNA kit (New England Biolabs) following manufacturer’s instructions^22^. For the tracheal aspirate samples described in Application 4, RNA was extracted using the Qiagen Allprep kit (Qiagen) following manufacturer’s instructions. Sequencing libraries were prepared using the NEBNext Ultra-II RNA kit (New England Biolabs) according to a previously described protocol^27^. Paired end 150 base pair illumina sequencing was performed on all samples using Illumina NextSeq 550 or NovaSeq 6000.

### AMR gene identification

We downloaded the tabular results from the AMR module and applied quality filters to ensure robust AMR gene identification. Specifically, for mNGS data, we required all AMR genes (from contig and read approaches) to have coverage breadth > 10% and for read mappings we additionally required > 5 reads mapping to the AMR gene. For WGS data, we required all AMR genes (from contig and read approaches), to have coverage breadth > 50% and additionally required > 5 reads mapping to the AMR gene for read results. Across all analyses, Nudged results were treated the same way as contig results. For studies with corresponding water controls, we applied the above filters to the water controls, and then removed AMR genes or gene families (depending on what was plotted) also found in water controls from experimental samples.

### AMR gene heatmaps

All plots were generated in R using Tidyverse^32^, patchwork^33^ and ComplexHeatmap^34^. While making the plots, we did an additional filtering to focus the analysis within the context of the use-case and limit the size of the plots for the paper. In particular, we included only CARD’s protein homolog and protein variation models (see https://github.com/arpcard/rgi), and included only medically relevant antibiotics drug classes by removing disinfecting agents and antiseptics, antibacterial free fatty acids, and aminocoumarin, diaminopyrimidine, elfamycin, fusidane, phosphonic acid, nucleoside, and pleuromutilin antibiotics. In Fig. 5B and Fig. 8, we also excluded efflux pumps to reduce plot size as efflux pumps tend to have ubiquitous functions in cellular processes.

Then, we applied a series of heuristics to make this structured data amenable to heatmap visualization. Given the nature of a heatmap visualization, each AMR annotation in each sample can have only one representing tile, so we plotted the result with the highest confidence. We considered AMR genes identified through the contig approach with Perfect or Strict cutoffs as higher confidence than those with the Nudged cutoff, which were then of higher confidence than AMR genes found by reads alone. Finally, given the challenges for gene attribution presented by homology between genes in the same gene family, we developed a systematic approach for collapsing the visualization to a single candidate per sample. For all figures except for Fig. 6, if in the same sample one AMR gene was found by the read approach and a different AMR gene from the same gene family was found by the contig approach, the first AMR gene was omitted and only the second AMR gene was plotted. The rationale for this prioritization stems from the fact that sometimes short reads alone cannot sufficiently distinguish between highly similar alleles or genes from the same gene family. Contigs, which typically provide greater sequence length are often of higher confidence. This approach should be considered on a per gene or per gene family basis, due to variability in the extent of sequence similarity within genes and gene families, and also be modified for specific use cases. For example in Fig. 6B, even though *mecR1* and *mecA* are from the same gene family, they do not have highly similar sequences and we did not apply this step.

### Species identification

For results from the CZ ID mNGS module, filters were again applied to ensure high-quality results. Specifically, for Fig. 3 and Fig. 7, which each focused on a single species, the NT rpM calculated by the mNGS module was used with no extra filtering. For Fig. 5 and Fig. 6A, which focused on species composition, the species detected by the mNGS module were filtered with: NT rpM > 10 and NR rpM > 10 to implement a minimal abundance requirement for taxonomic identification, NT alignment length > 50 to ensure alignment specificity and NT Z-score > 2 using a background model calculated with the corresponding study-specific water samples to ensure significance of taxa above levels of possible background contamination. Finally, for Fig. 6B, which had low read coverage, abundance filters were omitted and only the significance filter of NT Z-score > 2 was applied, using a background model calculated with the corresponding water samples.

### SNP distance analysis

Host-filtered reads were downloaded from the CZ ID mNGS module. SNP distance were calculated with SKA2 0.3.2^23^ using ska build--min-count 4--threads 4--min-qual 20 -k 31--qual-filter strict and ska distance--filter-ambiguous. The heatmap plot was generated with ComplexHeatmap^34^

### Data and code availability

All raw microbial sequencing data supporting the conclusions of this article are available via NCBI’s Sequence Read Archive under BioProjects PRJNA544865, PRJNA1086943, PRJNA450137 and PRJNA672704. For previously unpublished datasets, non-host FASTQ files generated by CZ ID mNGS module were submitted to SRA under NCBI Bioproject Accession: PRJNA1086943. We obtained raw FASTQ files from previous studies^22,25–29^, either from the authors or public repositories, and uploaded them to the CZ ID pipeline (https://czid.org/) under an openly accessible manuscript-specific project called “AMR example applications” to be processed through both the AMR module and the mNGS module (the project can be accessed at https://czid.org/home?project_id=5929 after logging in). CZ ID workflow code can be found in https://github.com/chanzuckerberg/czid-workflows/. Additional code for data filtering and plotting can be found in https://github.com/chanzuckerberg/czid-amr-manuscript-2024. The following software versions were used for this manuscript: CZ ID mNGS workflow version 8.2.5, CZ ID AMR workflow version 1.4.2 based on CARD RGI version 6.0.3, CARD database versions 3.2.6 and the CARD Resistomes & Variants database: 4.0.0. SK2 version 0.3.2.

## Authors’ contributions

KK and CL conceived of and designed the work. DL carried out data analysis with valuable inputs and guidance from KK, CL, VC and AG. ESG collected and sequenced all samples in Application 2. The CZ ID team (NB, XB, KR, KE, EF, OH, EH, AEJ, RL, SM, LR, JT, OV) built the AMR module. PMM collected and sequenced all samples in Application 4. AJP provided the data for Application 5. ARR, BPA, AGM provided expert input on the project. CL supervised the work. DL, KK and CL drafted the manuscript with inputs from all coauthors.

## Acknowledgements

We acknowledge the contributions of the whole CZI Infectious Disease development team: Robert Aboukhalil, Kami Bankston, Neha Chourasia, Jerry Fu, Julie Han, Francisco Loo, Todd Morse, Juan Caballero Perez, David Ruiz, Vincent Selhorst-Jones and Kevin Wang.

## Supplementary Materials

**Figure S1.**
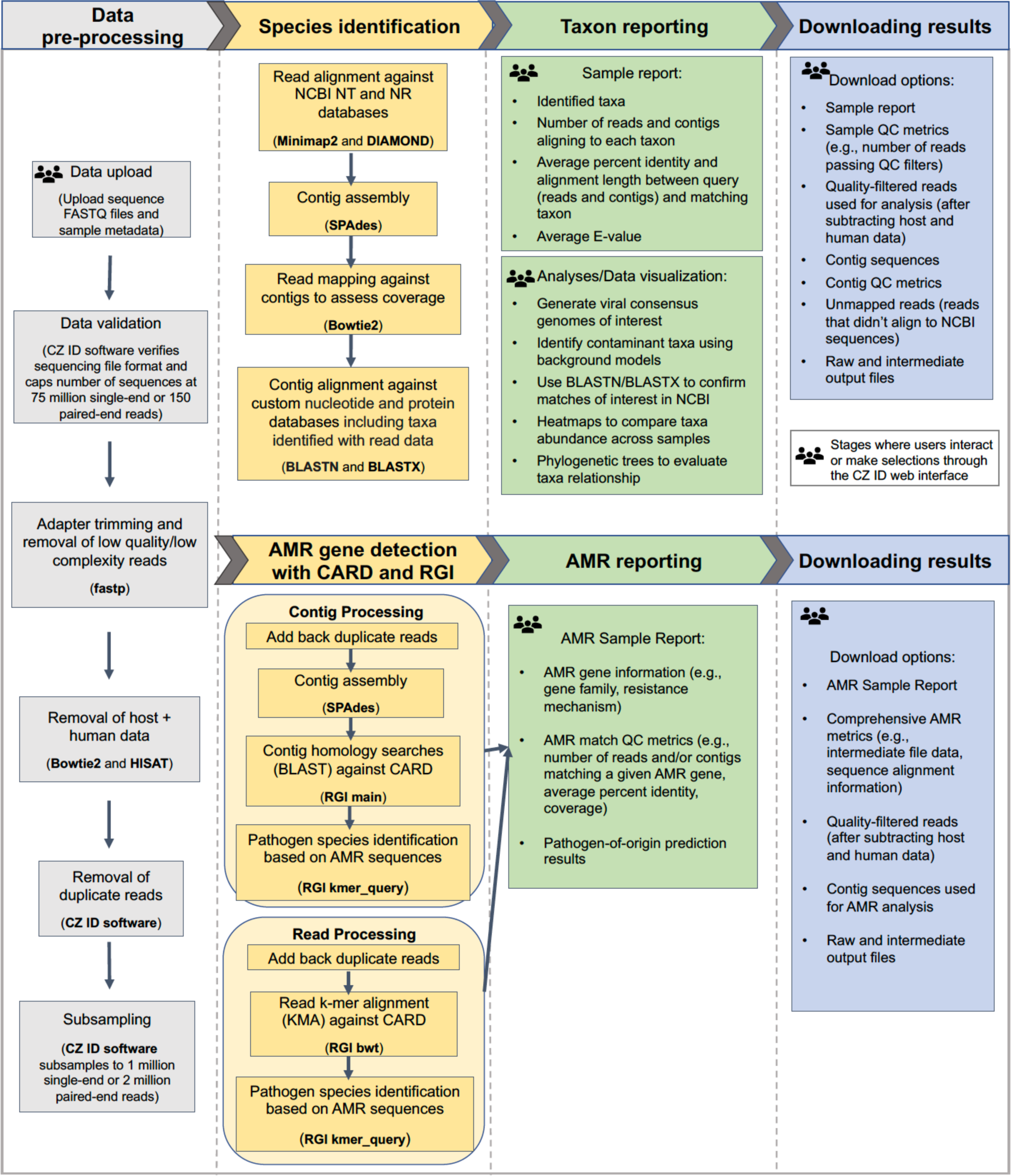
Detailed flow diagram highlighting the integrated AMR and mNGS modules within the CZ ID pipeline.

